# A key role of the hippocampal P3 in the attentional blink

**DOI:** 10.1101/2022.08.08.502473

**Authors:** M Derner, TP Reber, J Faber, R Surges, F Mormann, J Fell

## Abstract

The attentional blink (AB) refers to an impaired identification of target stimuli (T2), which are presented shortly after a prior target (T1) within a rapid serial visual presentation (RSVP) stream. It has been suggested that the AB is related to a failed transfer of T2 into working memory and that hippocampus (HC) and entorhinal (EC) cortex are regions crucial for this transfer. Since the event-related P3 component has been linked to inhibitory processes, we hypothesized that the hippocampal P3 elicited by T1 may impact on T2 processing within HC and EC. To test this hypothesis, we reanalyzed microwire data from 21 patients, who performed an RSVP task, during intracranial recordings for epilepsy surgery assessment (Reber et al., 2017). We identified T1-related hippocampal P3 components in the local field potentials (LFPs) and determined the temporal onset of T2 processing in HC/EC based on single-unit response onset activity. In accordance with our hypothesis, T1-related single-trial P3 amplitudes at the onset of T2 processing were clearly larger for unseen compared to seen T2-stimuli. Moreover, increased T1-related single-trial P3 peak latencies were found for T2[unseen] versus T2[seen] trials in case of lags 1 to 3, which was in line with our predictions. In conclusion, our findings support inhibition models of the AB and indicate that the hippocampal P3 elicited by T1 plays a central role in the AB.

## Introduction

Human visual attention has peculiar temporal limitations. The attentional blink (AB) refers to a transient impairment in the perception of visual stimuli, which are presented in rapid succession (Raymond et al., 1992). More precisely, the ability to identify and report a target stimulus (T2) is reduced when it appears with a short delay (typically 150-500 ms) after a prior target (T1). While numerous theories have been proposed to explain this phenomenon, a controversial debate is still ongoing (Dux and Marois, 2009; Snir and Yeshurun, 2017). An undisputed mechanistic account of the AB based on neurophysiological findings is yet missing.

Inhibition models have proposed that the AB results from a suppressive mechanism inhibiting the processing of stimuli occurring after target T1 (Raymond et al., 1992; Olivers et al., 2007). The event-related P3 component is observed in target-detection tasks (Donchin, 1981; Picton, 1992) and has been linked to inhibitory processes (Elbert and Rockstroh, 1987; Polich, 2007). The latency window of the P3 (typically 200-700ms) is well in line with the idea that the P3 elicited by T1 interferes with T2 processing. Therefore, a central role of the T1-related P3 in the AB has been proposed (McArthur et al., 1999; Fell et al., 2002). Indeed, based on surface recordings moderate associations of the T1-related P3 with the AB have been reported (e.g. Sergent et al., 2005). However, unambiguous evidence for a key role of the T1-related P3 in the AB has been lacking.

When T2-stimuli are not seen, early T2-related sensory processing appears to be largely intact, while the T2-related P3 is absent (Zivony and Lamy, 2022). Since the P3 has been related to conscious perception and working memory updating (Donchin, 1981; Polich, 2007), this may indicate a failure to transfer T2-stimuli into working memory. It has been suggested that the hippocampus (HC) is a major network hub for working memory processing (Fell and Axmacher, 2011; Kaminski et al., 2017; Kornblith et al., 2017) and that the entorhinal cortex (EC) represents its gateway (Fernández and Tendolkar, 2006). Based on human single-neuron data, it indeed has been shown that T2-related hippocampal and entorhinal population responses are markedly reduced for unseen versus seen T2-stimuli (Reber et al., 2017). Therefore, we hypothesized that the T1-related mediotemporal lobe (MTL)-P3, which is generated within the hippocampus (Halgren et al., 1980; Grunwald et al., 1999), is a crucial factor in the AB due to its impact on hippocampal/entorhinal processing of T2.

To investigate this hypothesis, we re-analyzed AB data recorded from 21 epilepsy patients undergoing invasive seizure monitoring in preparation for resective neurosurgery (Reber et al., 2017). In these patients mediotemporal depth electrodes and microwires had been implanted for chronic seizure monitoring. During 40 experimental sessions patients performed a rapid serial visual presentation (RSVP) task using images as stimuli (Figure 1A). These images were individually determined in a preceding screening session based on selective mediotemporal single-neuron responses. Behavioral data (Figure 1B) showed a pronounced reduction of target detection for those T2-stimuli, which were presented 150 ms (lag 0) or 300 ms (lag 1) after T1. To test the above hypothesis, we asked whether T1-related P3 amplitudes at the onset of hippocampal/entorhinal T2 processing allow to predict whether T2-stimuli are consciously perceived. The information displayed is concordant with the information displayed in figure 1 (A,B) of Reber et al. (2017).

**Figure 1:**
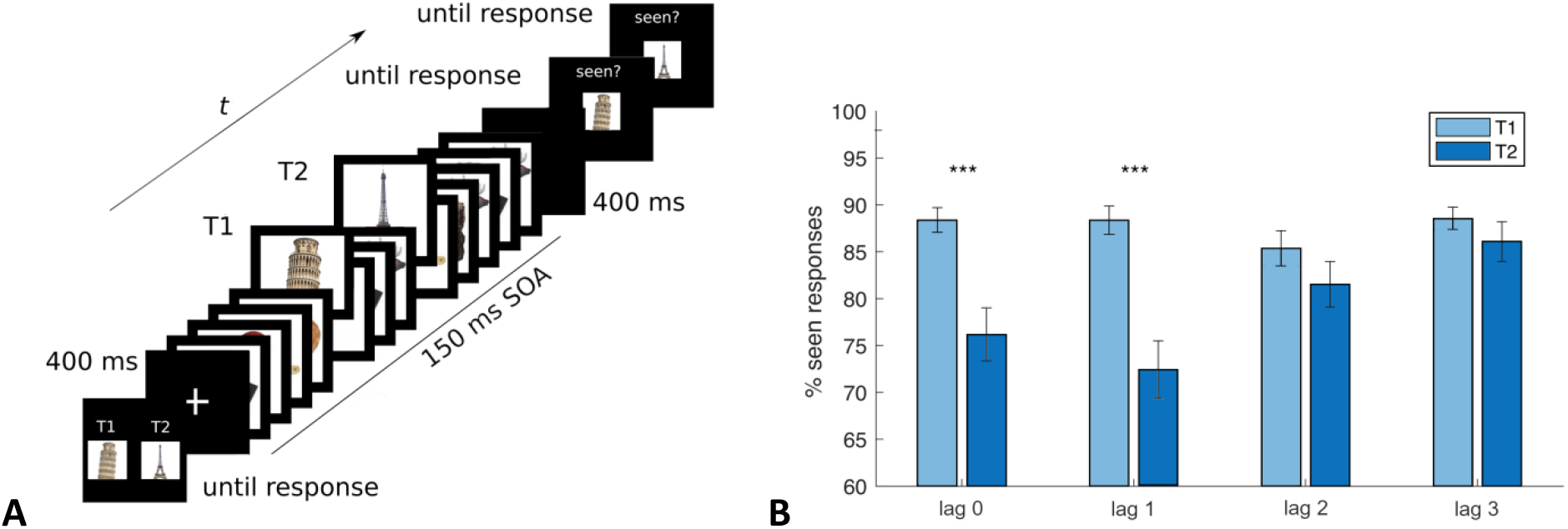
Experimental paradigm and behavioral results. (A) The sequence of events in an exemplary trial is shown from bottom left to top right. Eight subject-specific stimuli were chosen prior to the main experiment based on selective single-neuron responses in a preceding screening session. Subjects were asked to watch for two of these eight stimuli among 14 images presented in a rapid serial visual presentation (RSVP) sequence. The target stimulus that appeared first in the sequence is referred to as “T1” and the one that appeared second is referred to as ‘‘T2’’. The lag between T1 and T2 images varied from 0 to 3 (3 in the trial shown). The stimulus onset asynchrony (SOA) was usually 150 ms. After the RSVP stream, participants indicated with button presses whether they had seen T1 and T2 or not (two separate queries). Trials were classified accordingly into T1/T2 seen and T1/T2 unseen. (B) Average percentages of seen T1 and T2 images. Asterisks denote significant differences between T1 and T2 (post-hoc pairwise T-tests after significant target x lag interaction in 2 × 4 repeated measures ANOVA; lag 0, lag 1: p < 0.0001); error bars depict standard errors of the mean. Behavioral results indicate that T2-stimuli were less often reported to be seen than T1-stimuli for lag 0 (150 ms after T1) and lag 1 (300 ms after T1).

## Results

In a first step, we estimated the time range of the onset of T2 processing within HC/EC based on examination of single-unit response onset latencies. For this purpose, we determined the firing latencies of stimulus-responsive units (see Reber et al., 2017) in HC/EC (n=26) selectively responding to T2-stimuli for instances when T2-stimuli were seen (Figure 2). The median of T2[seen]-related firing latencies across stimulus-responsive HC/EC units was 308.2 ms, and the 25%- and 75%-quartiles were 240.7 ms and 391.7 ms, respectively.

**Figure 2:**
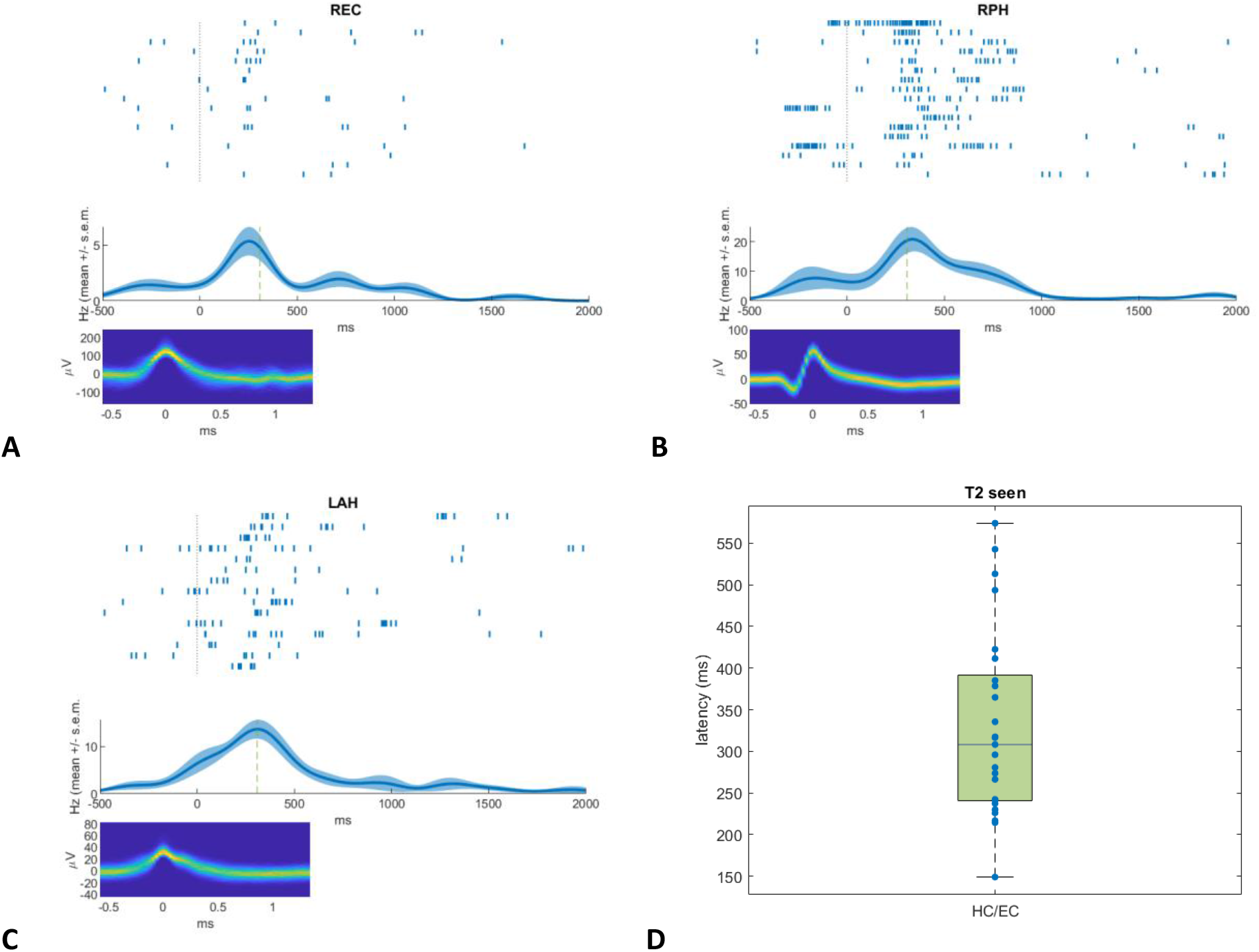
Examples of selective single-neuron responses and response latency of HC/EC neurons to seen T2-stimuli. (A-C) Three example units selectively responding to subject-specific T2-stimuli. Top: Raster plots of observed spike times relative to stimulus onset of T2 (vertical dotted line). Middle: Mean instantaneous firing rates (Hz). Zero on the x-axis denotes stimulus onset. Vertical dashed lines mark mean response latencies to T2[seen]. Bottom: Density plots of all spike waveforms. The plots show 2-dimensional histograms of spike voltages over time. The color code depicts the percentage of spikes (denominator: all spikes recorded for this unit) with the specified voltage at the given time point. REC, right entorhinal cortex; RPH, right posterior hippocampus; LAH, left anterior hippocampus. (D) Boxplot of firing latencies of n=26 stimulus-responsive units in hippocampus (HC) and entorhinal cortex (EC) responding to seen T2-stimuli. Blue dots mark the median response latency to T2[seen] stimuli in each unit.

In a second step, we identified T1-related hippocampal P3 components in the local field potentials (LFPs) recorded with the microwires. P3 components were visually scrutinized in accordance with previous reports based on intracranial electroencephalogram recordings (Halgren et al., 1980; Grunwald et al., 1999; Fell et al., 2005). More specifically, we searched for pronounced components peaking between 300 and 600 ms and clearly protruding from background activity. Because of the referencing scheme (see Materials and Methods) P3 identification was performed independent of polarity. A hippocampal P3 could be detected in 16 of 21 patients and 28 of 40 sessions (peak latency (average ± s.e.m.): 450.9 ± 8.5 ms; absolute peak amplitude: 27.9 ± 3.1 µV). In seven patients and 12 sessions, P3 components were identified in both hemispheres, and in nine patients and 16 sessions in one hemisphere. For each of these sessions and hemispheres, we chose the hippocampal channel showing the most pronounced P3 resulting in 40 cases overall. Finally, for each of these cases the microwire exhibiting the largest absolute P3 peak was selected (Figure 3).

**Figure 3:**
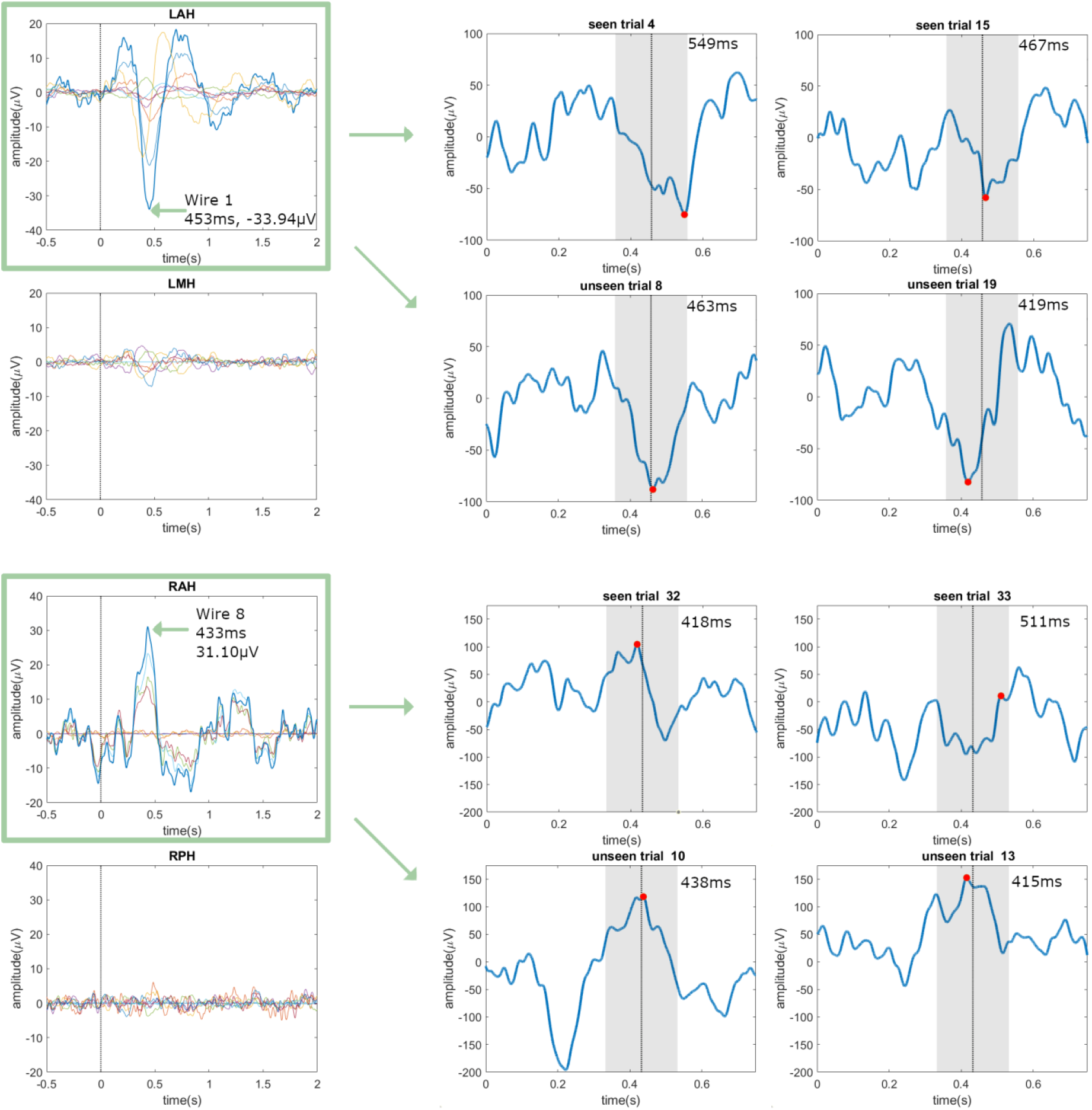
Selection of loci/wires exhibiting T1-related P3 components and single-trial analysis. Left column: Two examples for selection of the locus/wire with the most prominent T1-related P3 component (each wire in a different color). Mediotemporal P3 components in local field potentials were visually identified. They were assumed to peak between 300 and 600 ms and to be clearly distinguishable from background activity. Because of the referencing scheme the polarity of P3 components could either be positive or negative. For each session and brain hemisphere, the hippocampal channel with the most pronounced P3 was chosen (here shown: RAH (right anterior hippocampus) and LAH (left anterior hippocampus)). Finally, for each of these loci the microwire exhibiting the largest absolute P3 peak was selected (here shown: wire 8 for RAH and wire 1 for LAH; peak latencies and amplitudes are listed). The vertical lines mark the onset of stimulus T1; green boxes and arrows the selected channels and wires. Right column: Extraction of single-trial P3 peak latencies for four exemplary trials, each categorized as T2[seen] or T2[unseen] trial. Single-trial P3 peaks are defined as the maximum/minimum (according to the P3 polarity) amplitudes within +/-100ms around the case-specific average P3 peak latency. Vertical lines mark the latencies of the average P3 peaks, grey areas the +/-100ms intervals and red dots the single-trial P3 peaks. Single-trial P3 peak latencies are listed in the upper right corners.

As the central analysis, we performed a single-trial evaluation of T1-related LFPs for the 40 selected microwires (i.e. cases). LFP amplitudes were extracted at the time point of the median of T2[seen]-related HC/EC firing latencies, factoring in the trial-specific lags between T1 and T2. For each case, single-trial amplitudes were multiplied with the polarity sign (i.e. +1/-1) of the T1-related P3. Across cases, averaged single-trial LFP amplitudes were significantly larger for T2[unseen] versus T2[seen] trials (9.76 ± 2.65 vs. -6.35 ± 1.99 µV; p = 0.00024, paired one-tailed T-test; Figure 4A). Within cases, single-trial LFP amplitudes were significantly increased for T2[unseen] versus T2[seen] trials in 14 of 40 cases (unpaired one-tailed T-tests, each p < 0.05). A binomial test indicated that this number is significantly above chance level (p = 4·10^−9^). Moreover, average LFP amplitudes were calculated for the time interval corresponding to the [25%-quartile; 75%-quartile] of T2[seen]-related HC/EC firing latencies. Again, averaged single-trial LFP amplitudes were significantly larger for T2[unseen] versus T2[seen] trials across cases (7.93 ± 2.17 vs. -6.33 ± 1.80 µV; p = 0.00018; Figure 4B). Furthermore, in 19 of 40 cases single-trial LFP amplitudes were significantly increased for T2[unseen] versus T2[seen] trials (binomial test, p = 9·10^−15^).

**Figure 4:**
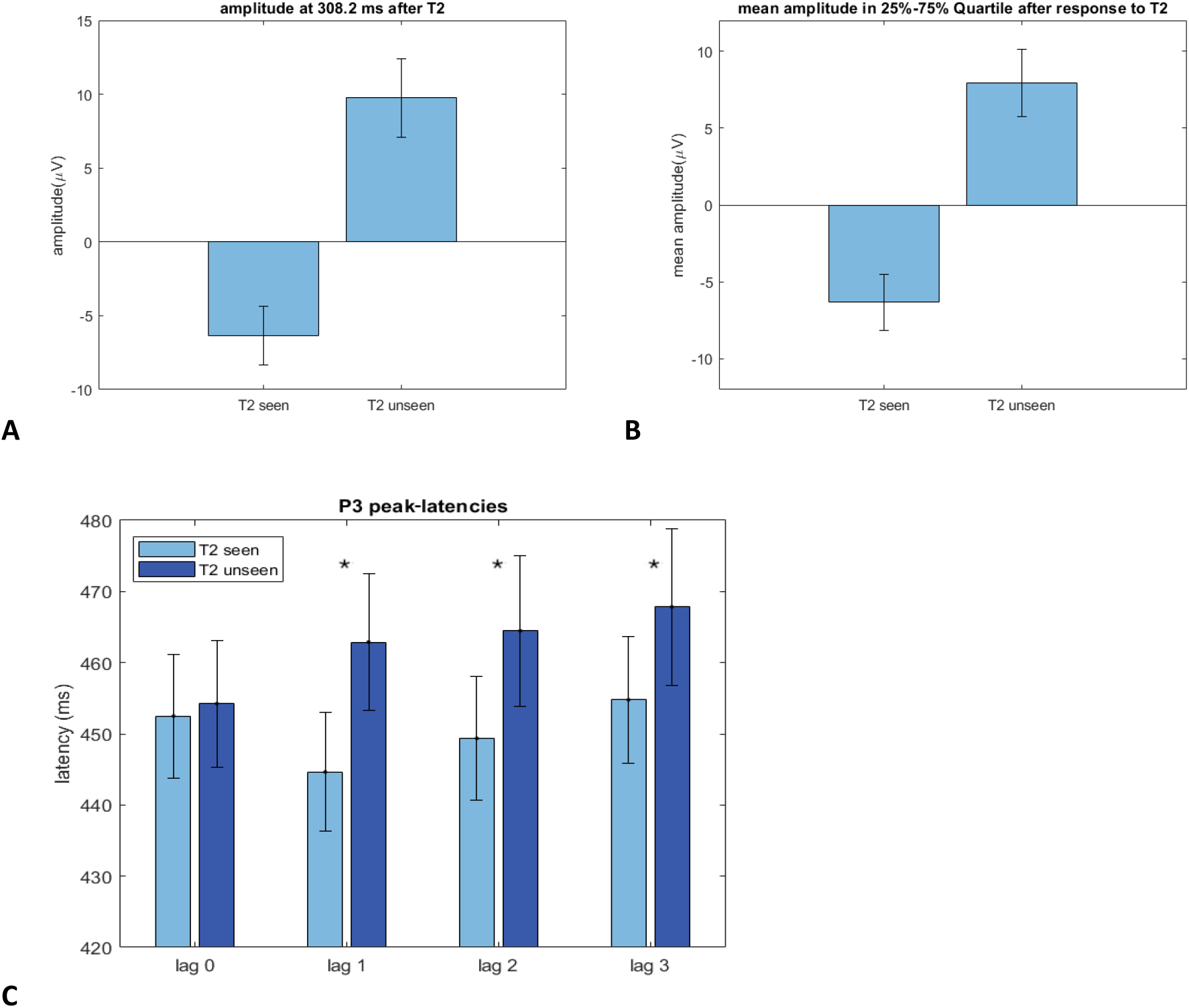
Amplitudes of single-trial LFPs for T2[seen] versus T2[unseen] trials and T1-related single-trial P3 peak-latencies. (A) Single-trial LFP amplitudes across cases at the time point of the median of T2[seen]-related firing latencies (mean and s.e.m. depicted). Single-trial amplitudes were multiplied with the polarity sign (i.e. +1 or -1) of the T1-related average P3 component. (B) Mean single-trial LFP amplitudes in the time interval corresponding to the [25% quartile; 75% quartile] of T2[seen]-related firing latencies. (C) Average T1-related single-trial P3 peak latencies for seen and unseen T2 images depending on the lag between T1 and T2. Asterisks denote significant differences between T2[seen] and T2[unseen] (two-tailed T-test for lag 0, one-tailed T-tests for lag 1, lag 2, lag 3); error bars depict standard errors of the mean.

We further asked, whether T1-related P3 peak latencies were different for unseen versus seen trials depending on the lag between T1 and T2. For lag 0 (150 ms), the peak of the T1-related P3 (average latency = 451 ms) typically occurred simultaneously to the onset of T2[seen]-related HC/EC firing (median latency = 308 ms). For lags 1 to 3 (300 to 600 ms), only P3 events with relatively long latencies might have an impact on T2-related HC/EC firing. Therefore, we hypothesized that single-trial P3 peak-latencies would be larger for T2[unseen] versus T2[seen] trials in case of lags 1 to 3, but would not differ in case of lag 0. To test this hypothesis, we evaluated single-trial P3 peak latencies, taking into account case-specific P3 polarities. More precisely, single-trial latencies of the maximum/minimum amplitudes within +/-100ms around the case-specific P3 peak latencies were extracted (provided P3 polarity was positive/negative, respectively; Figure 3). Indeed, P3 latencies were increased for T2[unseen] versus T2[seen] trials for lags 1, 2 and 3 (one-tailed T-tests across cases: p = 0.0001; p = 0.0038; p = 0.0273; Figure 4C). Moreover, P3 latencies did not differ between T2[unseen] and T2[seen] trials for lag 0 (two-tailed T-test: p = 0.65).

## Discussion

The present study reports the analysis of human LFP and action potential data recorded during an AB paradigm. Whether T2-stimuli were seen or unseen clearly depended on the amplitudes and latencies of the hippocampal P3 evoked by the T1-stimuli. These dependencies were in line with the idea that the hippocampal P3 impacts on T2-related processing within HC/EC and thereby may prevent conscious perception and transfer of T2-stimuli into working memory. More generally, these findings are in accordance with models suggesting that suppressive mechanisms inhibit the processing of stimuli presented after T1 (Raymond et al., 1992; Olivers et al., 2007), and with theories assuming a key role of the hippocampus in conscious perception (Behrendt, 2013; Berlucci and Marzi, 2019).

The P3 component has been related to a decreased excitability of cortical networks (Birbaumer et al., 1990; Elbert and Rockstroh, 1987). For instance, reaction times and evoked potential amplitudes in response to probe stimuli were prolonged (Rockstroh et al., 1992; Woodward et al., 1991) and startle reflexes were smaller (Schupp et al., 1994) after target stimuli eliciting a large P3. However, only moderate links of the T1-related P3 to the AB have been found based on surface recordings (McArthur et al., 1999; Sergent et al., 2005; Shapiro et al., 2006; Kranczioch et al., 2007). This suggests that the surface-recorded P3, which reflects contributions from several cortical generators (Soltani and Knight, 2000; Polich, 2007), may not be sensitive enough to capture the interference of T1-related processes with higher-order processing of T2-stimuli.

In conclusion, our data provide direct mechanistic evidence for the hypothesis that the hippocampal P3 elicited by T1-stimuli plays a central role in the AB. Our findings are in accordance with the theory that the hippocampal P3 interferes with processing of T2-stimuli within HC/EC at the level of conscious perception and transfer into working memory.

## Materials and Methods

### Participants

Recordings from 21 epilepsy patients (12 male; mean age: 37.9 ± 10.9 years) undergoing presurgical evaluation were re-analyzed (Reber et al. 2017). Mediotemporal depth electrodes and microwires had been implanted for chronic seizure monitoring and evaluation for epilepsy surgery. All patients gave informed written consent. The study conformed to the guidelines of the Medical Institutional Review Board at the University of Bonn (ethics votes Nr. 095/10 and 248/11).

### Experimental paradigm

A standard laptop running the Psychophysics Toolbox (Brainard, 1997) under MATLAB (MathWorks Inc.) was used for stimulus presentation. Subjects were asked to perform a rapid serial visual presentation (RSVP) task (Figure 1). The stimulus set for each of the 40 experimental sessions consisted of eight subject-specific images that were chosen based on selective mediotemporal single-neuron responses recorded in a preceding screening session (Kornblith et al., 2017). Participants were instructed to watch for two of these eight stimuli (T1 and T2) among 14 images presented in the RSVP sequence. At the beginning of each trial, a screen showing T1 and T2 was presented, and perception was confirmed with a button press. Then a fixation cross was presented for 400 ms, and thereafter the RSVP sequence of the 14 images started. The default stimulus onset asynchrony (SOA) was 150 ms (35 sessions), but was reduced to SOAs in the range of 100 to 135 ms (five sessions) in patients with only few unseen trials in their first experimental session. After the RSVP stream, there was a blank screen for 400 ms followed by two separate queries whether T1 and T2 had been seen or not.

Each session consisted of three runs of 72 trials each. The sequence of trials was randomized within each run. The eight response-eliciting images were chosen to be either T1 or T2 an equal number of times. To assess the false positive rate of seen reports, in 16 catch-trials per run either only T2 (eight trials) or T1 and T2 (eight trials) were omitted. The position of T1 and T2 in the sequence was set pseudorandomly with the constraints that T1 position ranged from 3rd to 5th, and that the lag between T1 and T2 varied from zero to three intervening images. The remaining 12 positions were pseudorandomly filled with the remaining six images with the constraint that identical images were not presented successively.

### Data recording

Recordings were obtained from a bundle of nine microwires (eight high-impedance recording electrodes, one low-impedance reference, AdTech, Racine, WI) protruding from the end of each depth electrode targeting hippocampus, entorhinal cortex, amygdala and parahippocampal cortex. Within the hippocampus, sections corresponding to the anterior, middle and posterior third were targeted: left/right anterior hippocampus (LAH/RAH: 21/20), left/right middle hippocampus (LMH/RMH: 17/13), left/right posterior hippocampus (LPH/RPH: 7/5). The differential signal from the microwires was amplified using a Neuralynx ATLAS system (Bozeman, MT), filtered between 0.1 and 9,000 Hz, and sampled at 32 kHz. These recordings were stored digitally for further analysis. The number of recording microwires per patient ranged from 32 to 96. Recording microwires were either referenced against one of the reference microwires or in a bipolar scheme, depending on signal quality. Signals were band-pass filtered between 300 and 3000 Hz. Spike detection and sorting was performed as described previously (Quiroga et al., 2004; Mormann et al., 2011).

### Identification of stimulus-responsive neurons (for detailed description see Reber et al. 2017)

Spike counts were obtained in overlapping 100-ms-bins within 0 to 1000 ms after stimulus onset and compared to the baseline window ranging from -400 to 0 ms for each presentation of an image. Based on the results of a Wilcoxon signed-rank test, the strength of the responses of each unit with regard to increased firing was quantified. Raster plots of unit responses with a p value < 0.001 were visually inspected by four experienced electrophysiologists and rated as valid responses or not. The following analyses focused on a subset of 26 stimulus-responsive neurons located in hippocampus and entorhinal cortex (see Reber et al., 2017).

### Computation of instantaneous firing rates

Z-scores of instantaneous firing rates were computed to compare neuronal firing across conditions. Instantaneous firing rates were calculated by trialwise convolution of spike trains with a Gaussian kernel (100 ms full width half maximum) and Z transformation of these signals with the mean and standard deviation in a baseline interval from -500 ms to 0 ms before stimulus onset across all target presentations (T1/T2). Normalized signals were averaged per unit and condition.

### Estimation of Response Latencies

Response latencies in a response period from 100 to 1000 ms after stimulus onset were estimated with a Poisson-burst detection algorithm (Hanes et al., 1995; Mormann et al., 2008) for units with a baseline firing rate above 2 Hz. For units with a lower baseline firing rate, firing latencies were estimated as the first spike time. The median of these response latencies across trials was calculated for the T2[seen] and T2[unseen] conditions for each unit. Only units where latency values could be determined for at least two trials per condition of interest (T2[seen], T2[unseen]) were included (25 of the 26 selected HC/EC units). For further analysis, the median firing latency (308.2 ms), as well as the 25% and 75% quartiles (240.7 ms; 391.7 ms) across stimulus-responsive units in hippocampus and entorhinal cortex responding to T2[seen] stimuli were calculated.

### Identification of P3 components

Analysis of local field potentials was performed in MATLAB using the FieldTrip toolbox (Oostenveld et al., 2011). Trials were segmented from -1000 ms to 2500 ms with regard to stimulus T1 onset and baseline-corrected with the baseline interval defined from -500 ms to 0 ms. Signals were bandpass-filtered from 1 to 30 Hz with a 2nd order Butterworth filter. To avoid edge effects, the resulting signals were cut to the interval from -500 ms to 2000 ms. Visual artifact rejection was performed and 4 % of all trials were discarded. Average local field potentials were calculated across all T1[seen] trials and hippocampal P3 components were visually identified. They were required to peak between 300 and 600 ms and to be clearly distinguishable from background activity based on visual inspection. Because of the referencing scheme the polarity of P3 components could either be positive or negative. For each session, the hemisphere-specific hippocampal channel (AH, MH or PH) showing the most pronounced P3 was chosen based on joint assessment of all microwires of each channel (Figure 3). A hippocampal P3 could be identified in 16 of 21 patients (LAH/RAH: 11/7; LMH/RMH: 3/1; LPH/RPH: 3/1) and 28 of 40 sessions (LAH/RAH: 17/10; LMH/RMH: 5/3; LPH/RPH: 4/1). Finally, for each of these channels the microwire exhibiting the largest absolute P3 peak was selected (Figure 3).

### Single-trial LFP amplitudes

For each of the 40 selected microwires and each trial, LFP amplitudes were extracted at the time point of the median of T2[seen]-related HC/EC firing latencies (308.2 ms) taking into account the trial-specific lags between T1 and T2. The single-trial amplitudes were then multiplied with the polarity sign (i.e. +1 or -1) of the T1-related average P3 component. Additionally, LFP amplitudes were extracted in the time interval corresponding to the [25% quartile; 75% quartile] of T2[seen]-related HC/EC firing latencies [240.7 ms; 391.7 ms]. These amplitudes were averaged across the time interval and likewise multiplied with the polarity sign. Across cases, the difference between averaged single-trial LFP amplitudes for T2 unseen versus seen trials was evaluated using a paired one-tailed T-test (hypothesis: amplitude [T2 unseen] > amplitude [T2 seen]). Within cases, single-trial LFP amplitudes for T2[unseen] versus T2[seen] trials were compared using unpaired one-tailed T-tests. Moreover, binomial tests with probability 0.05 (alpha level of 5%) were conducted to evaluate whether the number of cases with statistically significant increases of single-trial LFP amplitudes for T2[unseen] versus T2[seen] trials was higher than expected by chance.

### Single-trial P3 peak-latencies

Single-trial peak-latencies of T1-related P3 components were evaluated taking into account case-specific P3 polarities. In detail, single-trial P3 peak latencies were extracted as the time point of the maximum/minimum amplitude (according to the P3 polarity; positive: maximum, negative: minimum) within +/-100 ms around the case-specific average P3 peak latency (Figure 3). Single-trial peak latencies were categorized as related to T2[unseen] or T2[seen] trials and to T1/T2 lags of 0, 1, 2 or 3 for further analysis.

## Acknowledgements

This work was supported by the German Research Foundation (DFG MO 930/4-2, SPP 2205, SFB 1089).

## Data availability

In accordance with the ethics approval given by the ethics committee of the Medical Faculty of the University of Bonn and the guidelines of the German Research Foundation, pooled spiking data, local field potential data and program code will be made publicly available to researchers on a Github Online Repository. Further queries should be directly addressed to the corresponding author via email.

